# Factors affecting the juvenile departure in *Orientobdelloides siamensis* (Oka, 1917)

**DOI:** 10.1101/2024.04.17.589961

**Authors:** Poramad Trivalairat, Krittiya Trivalairat, Tashfia Raquib, Watchariya Purivirojkul

## Abstract

The departure of juveniles from parental care is an important period influenced by various factors. In the laboratory, 3-5 days after copulation, ten parent individuals of *Orientobdelloides siamensis* deposited approximately 361.6±37.79 eggs on the substrate and covered them until departure. The parents incubated their single-egg cocoons for around 7-9 days until the juveniles hatched. Subsequently, the newborns turned to attach to the ventral annuli of the parent using their caudal sucker. Between seven to eleven days after hatching, when the caudal sucker of juveniles expanded over the parent’s annuli, signaling their readiness to depart, they detached from beneath the parent vent to the substrate but continued to live beneath it. Finally, to determine the timing of juvenile departure, the insufficient space availability beneath the parental vent and yolk depletion around 14-21 days after hatching were analyzed. Through these morphological characteristics and behavior, this study indicated the interactions among these factors contributing to the mechanisms influencing juvenile departure in *O. siamensis*.

## Introduction

Glossiphoniidae Vaillant, 1890, is one of the freshwater proboscis leeches that distributed worldwide [1,2]. Distinguished by morphological characteristics, Glossiphoniidae also exhibits unique parental care behavior, for protecting and feeding of their young beneath the vent [1,2,3,4,5]. They are categorized into three subfamilies based on cocoon attachments: Glossiphoniinae (e.g., *Glossiphonia complanata* Linnaeus, 1758) attach cocoons to substrates, Haementeriinae (e.g., *Helobdella stagnalis* Linnaeus, 1758 and *H. triserialis* Blanchard, 1849) attach cocoons directly to the parent’s venter, and the monogeneric Theromyzinae (e.g., *Theromyzon tessulatum* Muller, 1774) exhibit a combination of these traits [4,6,7,8,9,10]. The placement of the cocoon on the substrate represents a primitive characteristic for this annelid worm, reminiscent of ancestors in polychetes, oligochaetes, and even primitive leeches (Infraclass Acanthobdellida) [2,4]. Furthermore, each species of glossiphoniid leeches has a distinct brooding period ranging from egg deposition to maturity, approximately 22 to 165 days [11,12]. However, a common trait among these juveniles is the departure of their parents after a single day. While some young leeches may experience yolk depletion and briefly separate from their parents, as observed in *G. complanata*, others, such as *H. stagnalis*, have a longer brooding period, from nourishing their young with a dietary meal to their juveniles until they fully mature [4]. Thus, aside from yolk storage, what other factors influence the parental departure of these young leeches?

Currently, there are four reported glossiphoniid leech species in Thailand: *Orientobdelloides siamensis* (Oka, 1917) (former *Placobdelliodes siamensis*), *O. sirikanchanae* (Trivalairat et al., 2019) (former *P. sirikanchanae*), *O. tridens* (Chiangkul et al., 2020a) (former *P. tridens*), and *Batracobdelloides bangkhenensis* Chiangkul et al., 2020b [12,13,14,15]. Most of these species are ectoparasites feeding on geoemydid terrapins, except for *B. bangkhenensis,* which preys on freshwater snails [2,5,12,13,14,15]. Among them, *O. siamensis* is a member of the Glossiphoniinae subfamily and the only species for which a complete life cycle and conditions have been reported. It boasts the shortest life cycle, lasting no more than a month from hatching to maturity, as well as the highest number of eggs per clutch and a large enough body size for observation with the naked eye [12]. To enhance our understanding of the departure of young leeches unaffected by parental nourishment, *O. siamensis* serves as a suitable model for parental departure examination. Therefore, we designed an examination to compare the caudal sucker diameter of juveniles and parent ventral annuli, body size between juveniles and parent, and the remaining yolk in the crop caeca associated with departure behaviours.

## Materials and methods

### Specimen preparation

Ten six-month-old specimens of *O. siamensis* (mature) were obtained from the laboratory of the Department of Zoology, Faculty of Science, Kasetsart University, Bangkok Province, Thailand (13° 50’ 53.6” N 100° 33’ 47.3” E), descendants from a previous study by Chiangkul et al. (2020). They were introduced into a large container measuring 57×32×33 cm³, which housed a Malayan snail-eating turtle (*Malayemys macrocephala* (Gray, 1859)), with water filled to two-thirds of the turtle’s height, for a few days to encourage feeding and copulation (16). After copulation, each specimen was isolated in a small container measuring 12×12×15 cm3 to facilitate egg deposition and raising of juveniles, and they were no longer provided with food (S1 Video). Water temperature in all containers was recorded every six hours (at 0000, 0600, 1200, and 1800 hours) daily. Throughout the observation period (1 February to 21 March 2024), the mean water temperature in the leech container was 29.1±2.4°C. Upon hatching, the number of juveniles from each parent was counted, and their crop caeca were observed daily for yolk storage until they left the parent. Yolk storage was assessed on a scale of 0 to 10 points: 10 points indicated full egg yolk in all crop caeca (comprising 6 anterior crop caeca and 4 post-caeca branches); the score was reduced by one when yolk in the crop caeca depleted from anterior to posterior, and 0 points indicated complete depletion of yolk in all crop caeca (Fig 1). Additionally, ten cocoons or juveniles from each parent, as well as all parents from the last day that the juveniles completely departed, were collected daily. These specimens were fixed in 4% glutaraldehyde for subsequent examination using scanning electron microscopy (SEM), and in 10% neutral buffered formalin for histological analysis.

**Figure 1.**
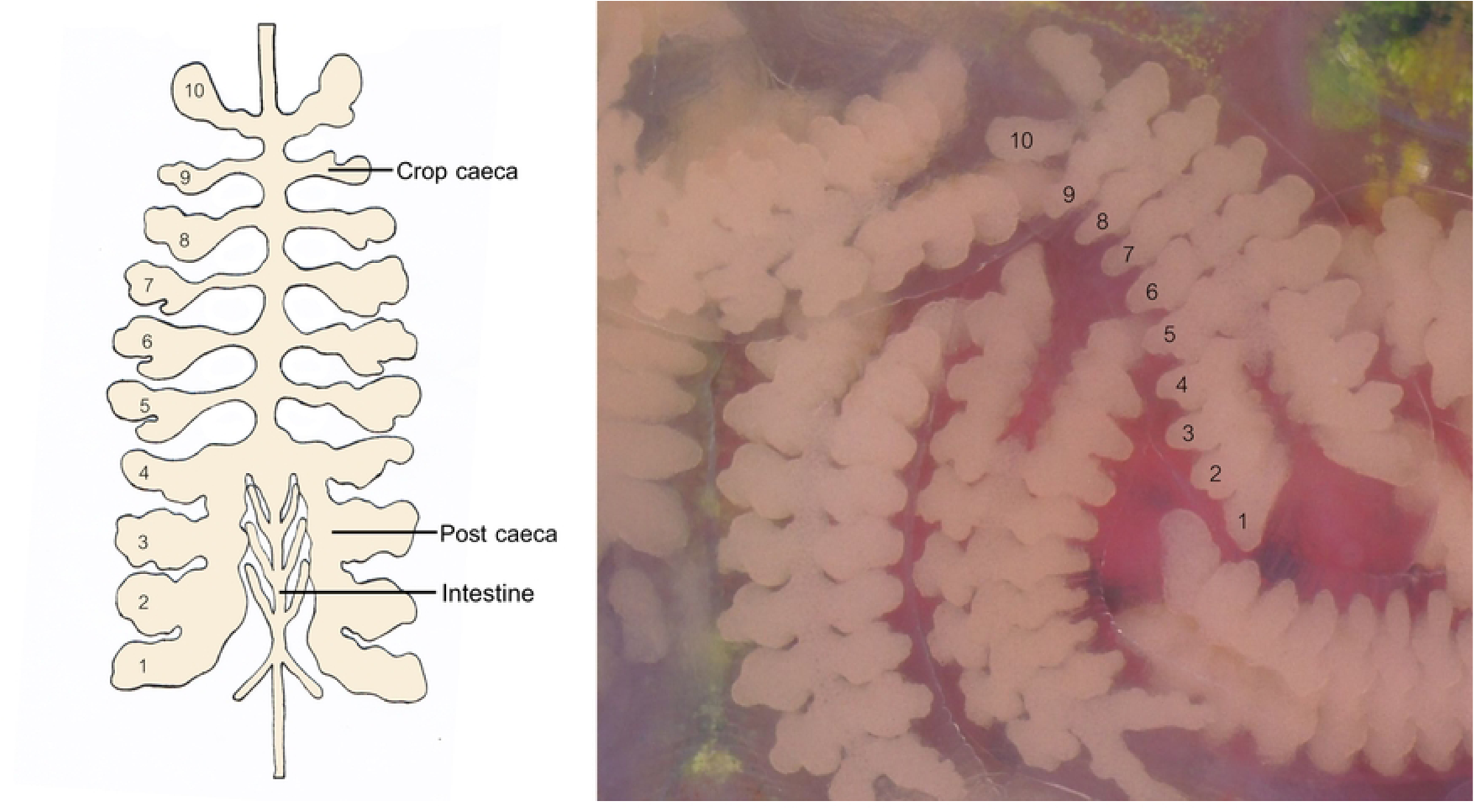
Yolk score assessment on a scale of 1 to 10 points, rang.

### Preparation of scanning electron microscopy technique

The specimens preserved in 4% glutaraldehyde were dehydrated in a series of ethanol solutions with two cycles spent in each concentration (70%, 80%, 95%, absolute ethanol; 1 hour for each concentration). This dehydration procedure was conducted using a critical point dryer (CPD; model HCP-2) until the specimens reached a critical dry point. The specimens were placed onto a stub using carbon tape and then sputter-coated with gold particles. All coated specimens were examined using a scanning electron microscope (SEM) to count the parent ventral annuli, measure the body length and width, including the length and width of the cocoon, as well as determine the diameter of the oral and caudal sucker of both parent and juvenile leeches. Additionally, this glossiphoniid leech was determined to have an oval shape, and the body area was calculated using the oval formula (π x (body width/2) x (body length/2)) for both parents and offspring. Furthermore, the capacity of offspring beneath the parent vent was calculated by multiplying three-fourths of the parent’s area by the offspring’s area on each day.

### Preparation of histological studies

The specimens were preserved in 10% NBF for 12 hours before dehydrating through a series of ethanol solution. They were cleared in xylene and embedded in paraffin blocks. Subsequently, the specimens were then sectioned into 5-mm-thick serial longitudinal sections using a rotary microtome. Following sectioning, the specimens were stained with hematoxylin and eosin (H&E) and mounted with Permount. The histological slides were examined under a light microscope to measure the anterior height (AH), posterior height (PH), and width (AW) for summing up the cross-sectional area (CSA) of each annulus (total of 76 annuli) (Fig 2). All photomicrographs were processed using Photoshop CS6.

**Figure 2.**
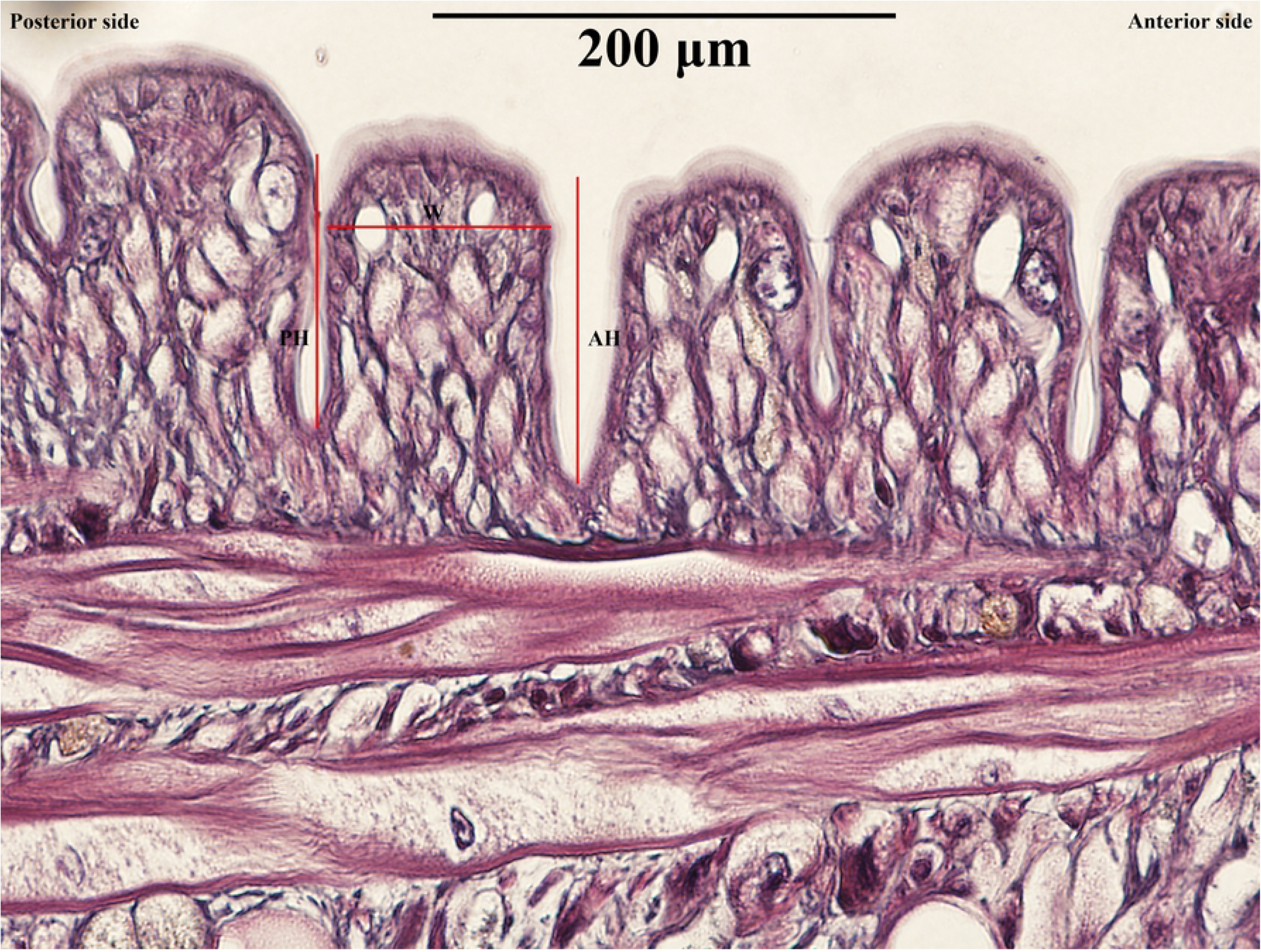
Measurements of annuli in Orientobdelloides siamensis.

### Ethic statement

This research was approved by the Institute of Animal Care and Use Committee at Kasetsart University under number ACKU66-SCI-016.

## Results

### Pre-hatching

Before the experiment, all ten one-year-old *O. siamensis* were starved for two weeks. On 17 February 2024, they were introduced into a large water container with a Malayan snail-eating turtle for feeding and stimulating a copulation (Fig 3). These leeches are active during the night. After two days, on 19 February 2024, all the parent leeches exhibited white creamy eggs inside their oviducts and turtle blood filling their crop caeca. At this point, they were transferred to separate small containers for isolation. Most of them deposited a clutch of cocoons between the third and fifth days following isolation (22 to 25 February 2024). These cocoons were a single-egg cocoon. The clutch sizes averaged approximately 361.6±37.79 eggs (range: 317–415 eggs, n = 10), with an average cocoon length of 455.75±36.53 μm (range: 363.28–564.97 μm, n = 100) and a cocoon width of 409.33±39.12 μm (range: 298.46–478.14 μm, n = 100) (Fig 4) (S1 Table). Ten cocoons were collected from each parent for analysis. Subsequently, the eggs turned a creamy brownish color during the fifth and sixth days after deposition (27 February to 2 March 2024) and hatched within a few days thereafter (29 February to 5 March 2024).

**Figure 3.**
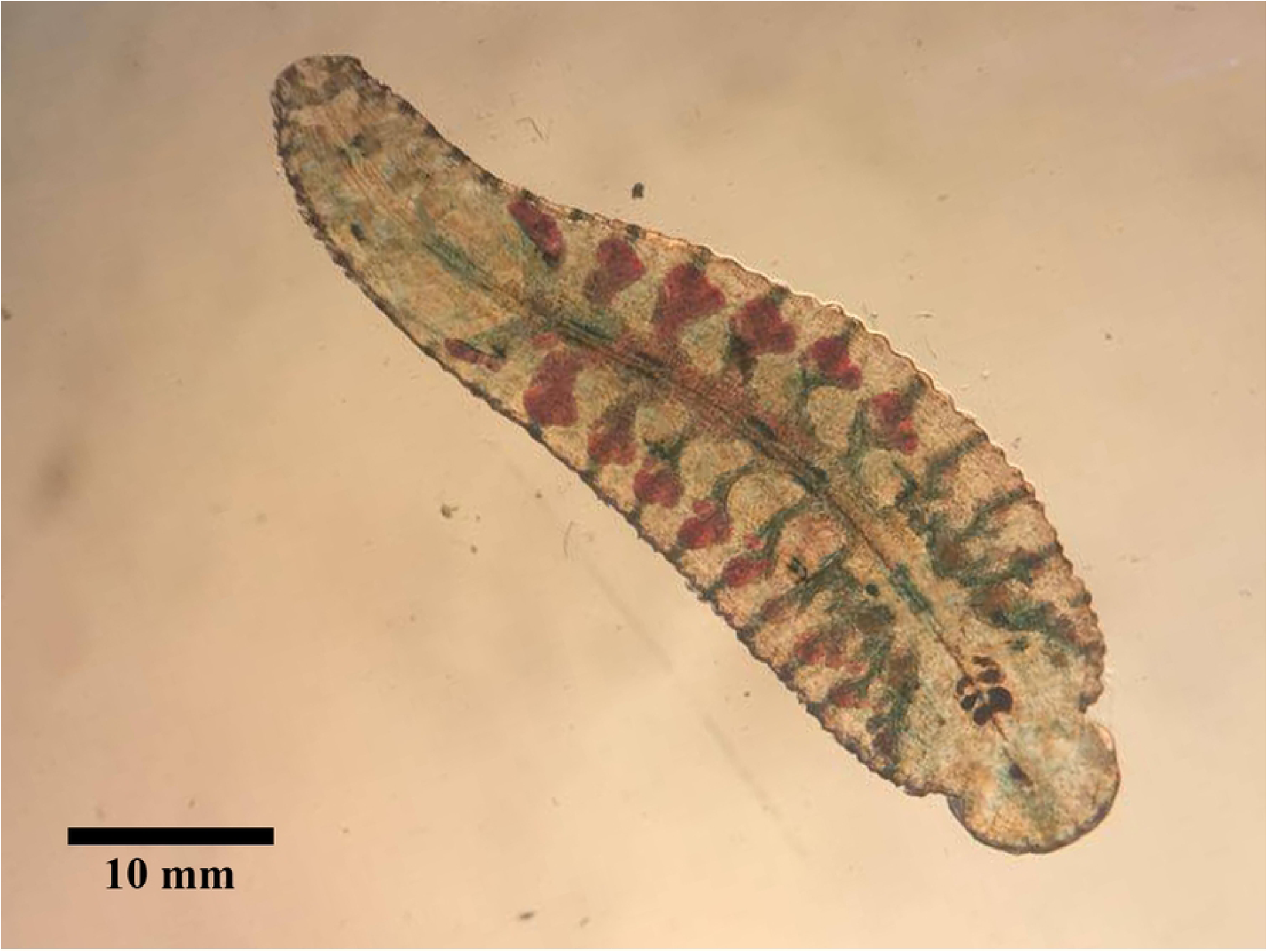
Mature Orientobdelloides siamensis exhibiting blood.

**Figure 4.**
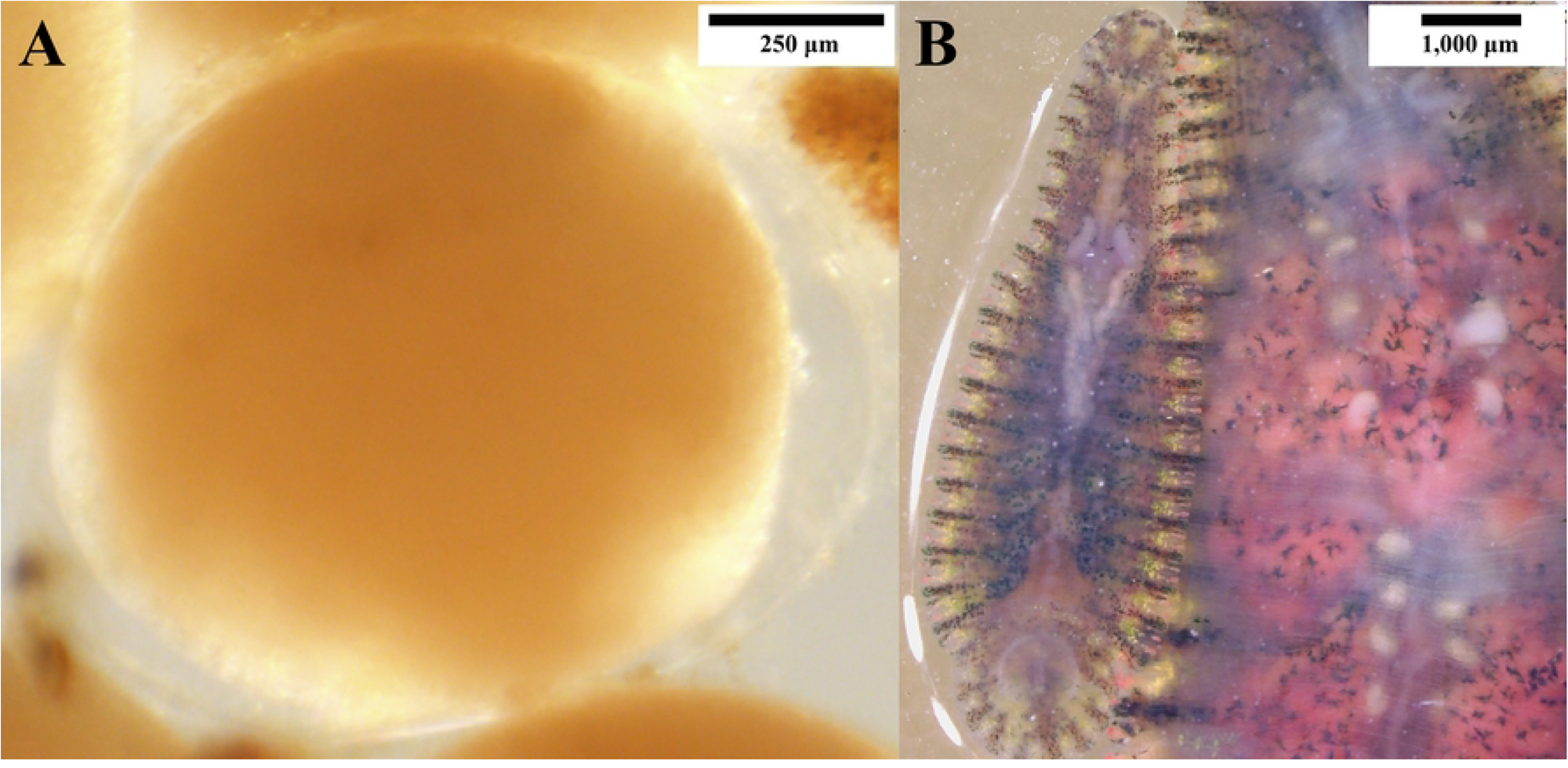
(A) Translucent cocoon containing an egg and (B) twenty.

### Post-hatching

The juvenile leeches inhabited beneath their parent’s vent until the second week post-hatching, undergoing a transition to attach to the substrate from the seventh to the eleventh day after hatching (7-11 March 2024), while remaining in close proximity to their parent. Throughout this transitional phase, the juveniles demonstrated a significant increase in body dimensions, with lengths spanning approximately 1,059.60-3,480.97 μm (1.88-6.18 times compared to the hatching day), widths ranging from 576.47-1,778.29 μm (1.54-4.74 times), OSD varying between 314.59-737.95 μm (1.79-4.20 times), and CSD ranging from 408.86-936.71 μm (1.66-3.79 times) (Fig 5).

**Figure 5.**
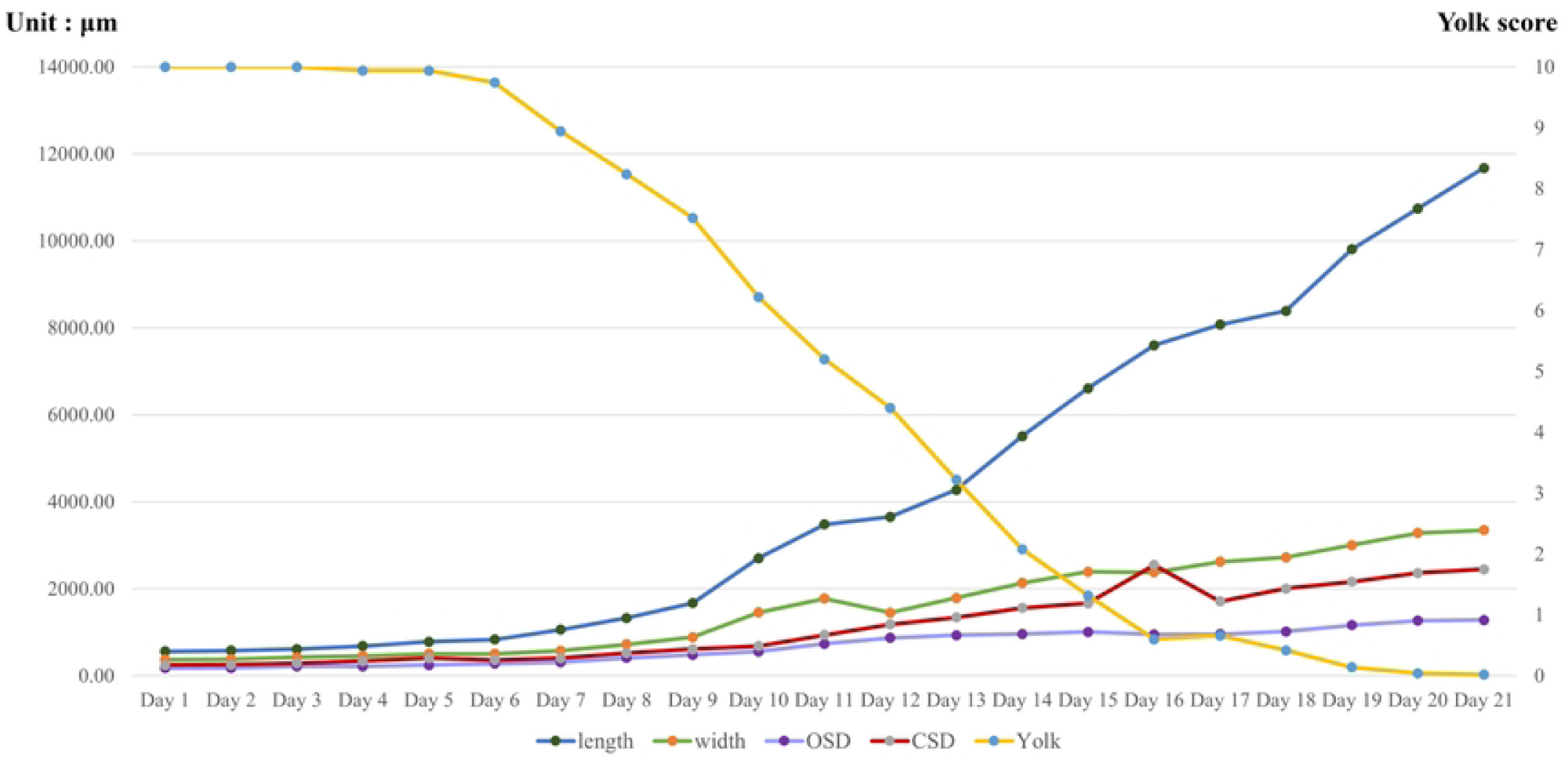
Growth rates of various characteristics, including body.

In comparison, the parental leeches displayed an average body length of roughly 53,802.04±5,632.71 μm (range: 49,232.57-61,045.56 μm), a width of 26,475.53±4,609.98 μm (range: 21,218.91-32,175.33 μm), OSD measuring 9,147.65±2,715.93 μm (range: 6,169.15-12,542.45 μm), and CSD of 10,874.05±2,222.81 μm (range: 8,905.47-13,418.16 μm), with a total of 76 annuli categorized into anterior and posterior sections. After the juveniles departed from their parents on the 21st day post-hatching (21 March 2024), the parents underwent fixation and histological processing to measure the total cross-sectional area (CSA) of each annulus by combining the widths and heights (anterior (AH) and posterior heights (PH)) of the annuli (as illustrated in S2 Table). Most anterior annuli exceeded a height of 100 μm, except for those around the gonopore (99.35-144.30 μm), corresponding to the genital pore areas (the male gonopore commonly found between the 20 and 21 annuli, varying between 20/21, 22/23, 25/26, and 26/27, and the female gonopore usually positioned between the 22nd and 23rd annuli, varying between 22/23, 24/25, 27/28, and 28/29). The posterior part, where juveniles distributed, exhibited an average CSA of approximately 432.90±68.31 μm (range: 280.36-611.98 μm), segmented into 261.77±67.08 μm in width (range: 127.95-383.86 μm), 81.25±27.63 μm in AH (range: 29.22-130.46 μm), and 89.87±30.56 μm in PH (range: 32.32-144.30 μm).

Seven days post-hatching (7 March 2024), the juveniles initially shifted their attachment from the vent to the substrate, and notably higher when their average CSD surpassed their posterior CSA on the eighth day (526.16±81.13 vs 432.90±68.31 μm), although the precise number of attachment transitions could not be determined (Fig 6). All juveniles completely transitioned to substrate attachment by the eleventh day post-hatching (11 March 2024), nearly doubling their CSD compared to three days earlier (936.71±162.51 μm, ratio 1.72) or 3.66 times compared to birth size (Fig 7).

**Figure 6.**
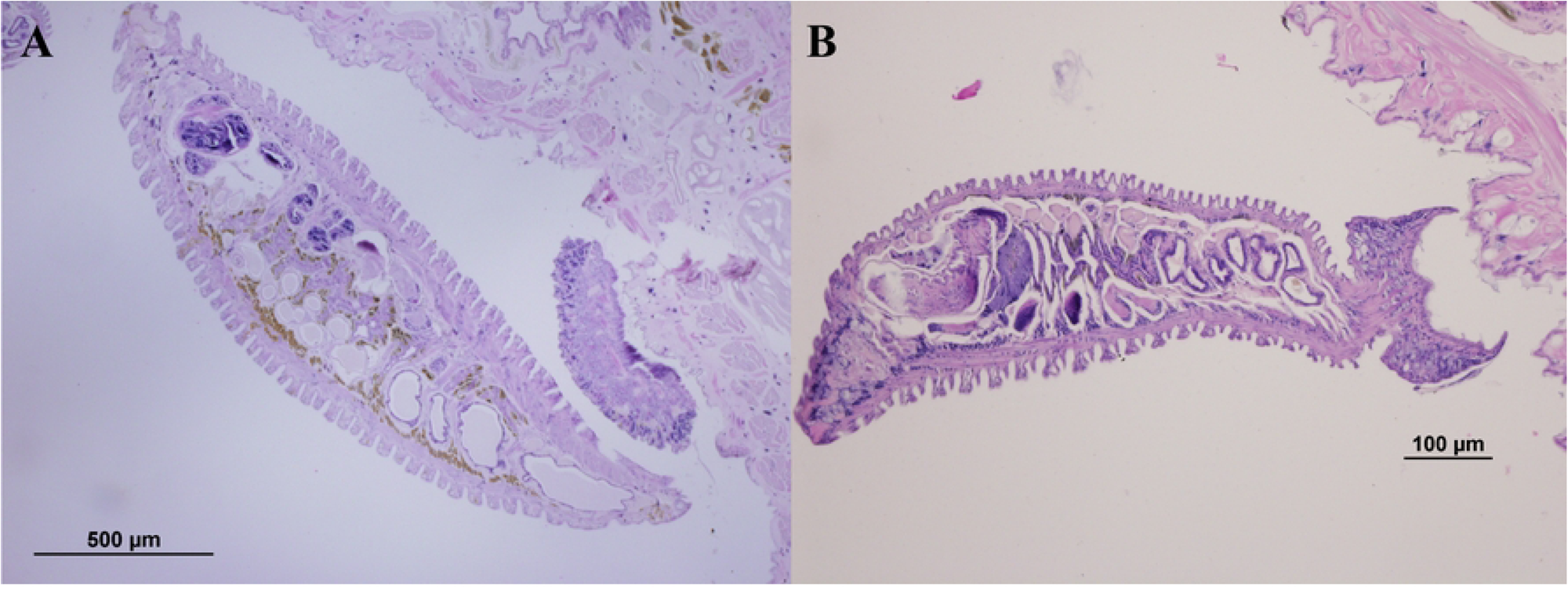
Attachment of Orientobdelloides siamensis juvenile by.

**Figure 7.**
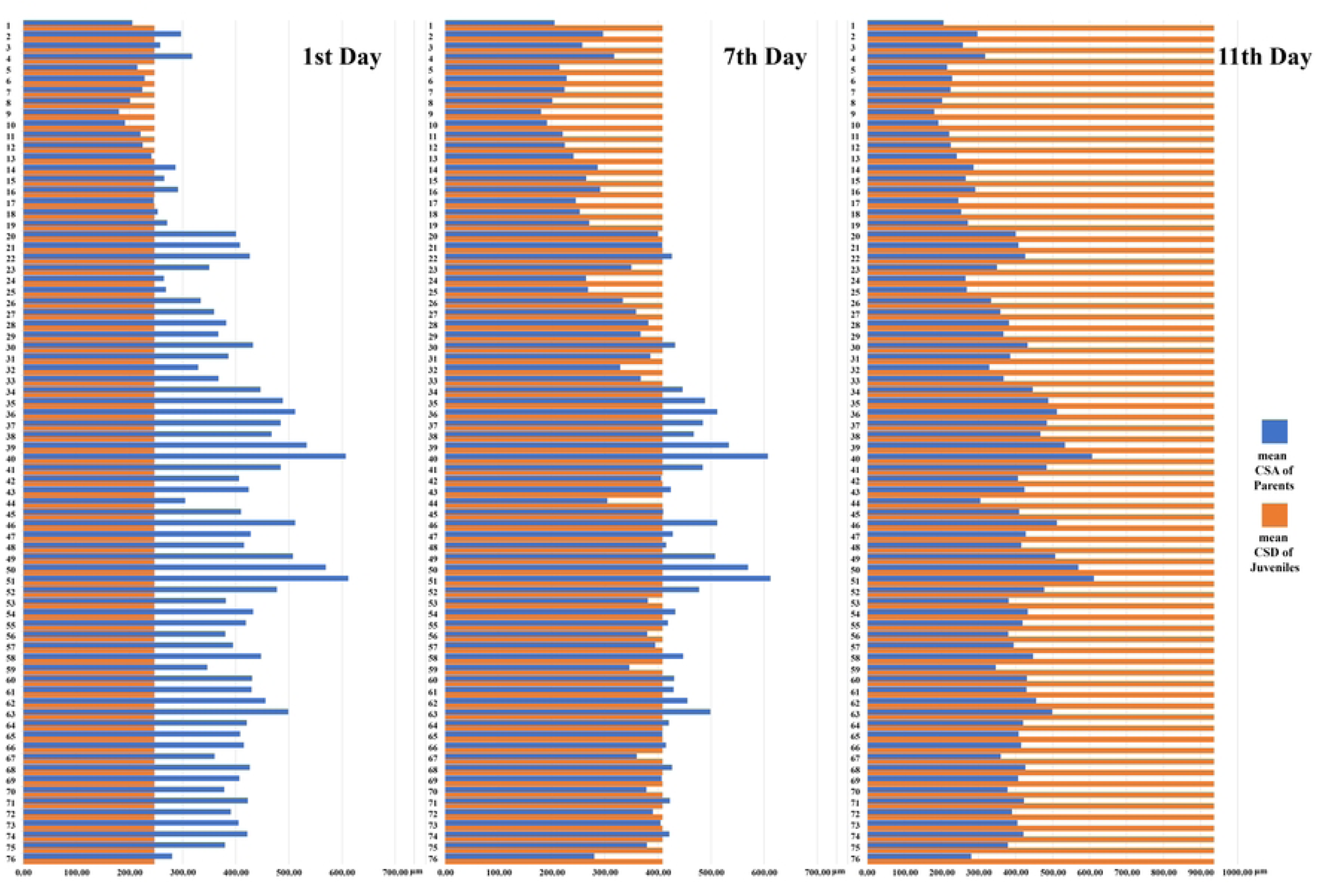
Comparison of parent annuli (annuli 1-76) and mean ca.

Additionally, on 10 March 2024, the average remaining yolk score decreased by almost half to 5.2, and four days later (14 March 2024), the first juveniles began to move away from its parent’s vent as their yolk reserves depleted. These 14-day-old juveniles measured 5,510.72±749.83 μm (range: 4,164.59-8,129.68 μm) in length, 2,133.33±291.91 μm (range: 1,542.29-2,935.32 μm) in width, 964.41±139.52 μm (range: 711.74-1,281.14 μm) in OSD, and 1,558.01±224.43 μm (range: 1,103.20-2,206.41 μm) in CSD. Subsequently, other juveniles that had depleted their yolk reserves also commenced moving away from their parents. The last day they cohabited with their parents was when they reached twenty-one days of age (21 March 2024). On this day, their average body size was approximately 11,676.25±1,764.47 μm in length (range: 7,603.06-16,988.09 μm), 3,351.13±691.35 μm in width (range: 1,841.69-5,059.99 μm), 1,283.55±375.59 μm in OSD (range: 607.70-2,651.80 μm), and 2,447.81±471.75 μm in CSD (range: 1,565.96-4,037.54 μm).

## Discussion

### Oviposition and Taxonomic Characterization

In the previous study from Chiangkul et al. (2020a), it was noted that *O. siamensis* (former *P. siamensis*) deposited cocoons and brood juveniles on the ventral surface as well as the behavior described in the subfamily Haementeriinae [12]. The previous observation was different from the description in Sawyer (1986) that classified this species under the subfamily Glossiphoniinae based on its cocoon deposition on substrates [2]. However, upon evaluation in this study, we found that *O. siamensis* deposited their tiny single-egg cocoons compared to their parent size on the substrate (455.75 vs. 53,802.04 μm for length, 409.33 vs. 26,475.53 μm for width) before the juvenile hatch and attach to their posterior vent (around three-fourths of the body). Thus, this behavior indicated that *O. siamensis* is the species that deposited on substrate and belongs to the subfamily Glossiphoniinae following Sawyer (1986), which currently encompasses a total of seven glossiphoniid genera including *Baicaloclepsis* Lukin and Epshtein, 1959, *Glossiphonia* Johnson, 1816, *Hemiclepsis* Vejdovsky, 1884, *Parabdella* Autrum, 1936, *Placobdella* Blanchard, 1893, *Torix* Blanchard, 1893, and *Orientobdelloides* [2,17,18,19,20,21,22,23]. Furthermore, the behavior observed in Gossiphoniinnae leeches represents a transitional stage between non-parental care seen in erpobdellid or hirudinid leeches (Arhynchobdellida) and developed parental care observed in theromyzinae leeches, which possess a brood pouch for carrying and protecting cocoons and juveniles [4,24,25,26,27]. Additionally, the parental care in glossiphoniid leeches culminates in the subfamily Haementeriinae, which exhibits developed feeding behavior towards their juveniles until they reach maturity.

### Affected departure factors

Typically, the caudal sucker of leeches plays a key role in adhering to the substrate through wet adhesive force (for smooth substrate) or contraction of muscle fibers (for rough substrate) [28]. Additionally, leeches used these adhesive properties in both their caudal and oral suckers, facilitating a distinctive mode of locomotion among hirudinea worms. In this study, the newly hatched *O. siamensis* transitioned their attachment from the substrate to their parent’s ventral surface by using their caudal sucker to adhere onto the annuli. Initially, the diameters of most caudal suckers (CSD) in newborns were smaller than the cross-sectional area (CSA) in the posterior ventral region of the parent (246.87 vs. 432.90 μm). This disparity in size between CSD and CSA facilitated their adherence onto the annuli. Despite the challenge posed by the rough surface of the parent’s annuli in gripping on, grip on the tip annuli for prior CSD expand and changing to grip on the whole annuli (as illustrated in Fig 5), the caudal sucker consisted of soft collagen tissues and highly ductile epidermal tissues, aiding in adhesion [28,29]. However, as the juveniles grew larger, their caudal sucker also increased in size until it surpassed the dimensions of the parent’s annuli. On the seventh day after hatching, some CSD measurements of the juveniles exceeded those of the parent’s annuli (408.86±52.27 μm, range: 305.41-551.35 μm), signifying the day when the juveniles transitioned their attachment back to the substrate. The last observed day of attachment to their parent was the eleventh day after hatching, with almost quadruple (3.66 times) in the size of the caudal sucker (936.71±162.51 μm, range: 696.20-1,376.58 μm). In theory, the larger size of the caudal sucker may provide leverage and a greater range of motion, potentially resulting in reduced force required for gripping compared to the smaller caudal sucker observed in newly hatched individuals [28,30,31]. Therefore, although the caudal sucker is indispensable for leech survival, the growth in its size signifies changes in juveniles, facilitating their first departure from the parent by reducing the adhesive force needed for attachment.

Moreover, upon hatching, the newly hatched juveniles of *O. siamensis* exhibited a substantial size difference compared to their parents. Their body length was nearly a hundred times smaller (95.51 times, 563.30 vs. 53,802.04 μm), while their body width, OSD, and CSD similarly decreased by factors of 70.50 times (375.53 vs. 26,475.53 μm), 52.04 times (175.78 vs. 9,147.65 μm), and 44.05 times (246.87 vs. 10,874.05 μm), respectively. Generally, the large juvenile comes with high energy acquisition from parents [32,33,34]. Some animals, including *O. siamensis*, might adapt to reduce the energy investment in offspring by decreasing the size but increasing the number instead, thereby enhancing their survival rate. However, these juveniles exhibited a rapid growth rate, emphasizing a 9.78-fold increase in length and a 5.68-fold increase in width when they initially departed on the 14th day compared to the hatching day (14 March 2024). They reached a remarkable 20.73-fold increase in length and an 8.92-fold increase in width on the last day of departure (21 March 2024).

Furthermore, examination of the oval-like body dimensions revealed the area beneath the parental vent, signifying available space for the juveniles. This area could accommodate all juveniles for a duration, based on an individual area of 166,205.72 sqμm on the hatching day. Remarkably, only three-fourths of the total body area (839,400,570.42 sqμm) was utilized, originating from the foremost attachment point (annuli 30), offering significant capacity for newly hatched juveniles, with a maximum of 5,050.37 individuals (maximum eggs per clutch in this study is 415). As they grow up, the juveniles could adjust by overlapping each other to conserve space, while their parents could expand their bodies to provide additional space. The results indicated that the space under the parental vent initially exceeded the capacity to accommodate all juveniles by the 10th day post-hatching (271.21 individuals), declining to 27.30 individuals by the 21st day (the last day of observed parental care). However, observations showed that juveniles commenced leaving the parent by the 14th day and had completely dispersed by the 21st day. Similar to other brooding animals, such as mouth-brooding fish, where body size plays a central role in the juvenile’s decision to depart from the mother’s mouth, this underscores the influence of the relationship between the space capacity in the natal habitat (under vent) and the size of the juveniles on their departure from the parent [35].

Lastly, the storage yolk, a significant determinant influencing the departure of offspring, plays a crucial role in leech development. Leech yolk, rich in essential nutrients such as vitellin and vitellogenin proteins, serves as a primary source of sustenance for juvenile leeches prior to independent feeding [36,37]. Among glossiphoniid leeches, *O. siamensis* stands out for its substantial investment in oogenesis, evident in the copious yolk reserves it allocates to its eggs. This species exhibits prolific reproduction, with clutches containing an average of 361.6±37.79 eggs (range: 317–415 eggs) observed in this study. In comparison to related species such as *Orientobdelloides* and *Helobdella*, which typically lay clutches comprising 50-200 eggs, O. siamensis demonstrates a remarkable two to eightfold increase in fecundity [4,14,24,25,26,38,39]. In this study, storage yolk deposition was visualized through transparent crop caeca, a feature exclusive to juvenile leeches. Initially, yolk presence was scored at 10, denoting complete occupancy across all branches (six in anterior caeca and four in post-caeca). Over the course of ten days, juveniles depleted approximately half of their yolk reserves (score 5.2), nearing complete exhaustion by the 21st day, coinciding with their departure. Notably, while yolk depletion paralleled body growth (as illustrated in Fig 3), *O. siamensis* diverges from typical reproductive strategies by extending parental investment beyond oogenesis to encompass parental care [40]. Despite lacking feeding behavior akin to helobdellid leeches, *O. siamensis* exhibits a unique reproductive strategy, parasitizing turtle shells for egg deposition, thus providing offspring with easier access to a blood meal compared to helobdellid counterparts [4,25,41,42,43]. Consequently, yolk provision represents a crucial parental investment in the initial growth and development of offspring, ensuring their survival prior to independent feeding. Moreover, the selection of suitable egg deposition sites enhances offspring survival by facilitating continued access to nourishment post-yolk depletion.

In summary, the departure of juvenile *O. siamensis* during the brooding period is governed by multiple factors. Initially, after hatching, the newborn juveniles undergo an attachment stage beneath the parent vent, which occurs around days 7-11 (Fig 8). This stage is characterized by their adherence to the parent’s ventral surface, facilitated by the caudal sucker. Following the attachment stage, the de-attachment stage begins around days 7-11, lasting approximately 10-14 days before the departure stage. During this phase, the expansion of the caudal sucker enables detachment, signaling their readiness to depart. Subsequently, the departure stage ensues, marked by yolk depletion and the accommodation of juveniles as they mature. Additionally, parental care ensures survival, while the deposition of eggs on turtle shells provides access to nourishment. Collectively, these elements, including caudal sucker development, yolk provision, and available space, play pivotal roles in shaping the departure and survival of offspring.

**Figure 8.**
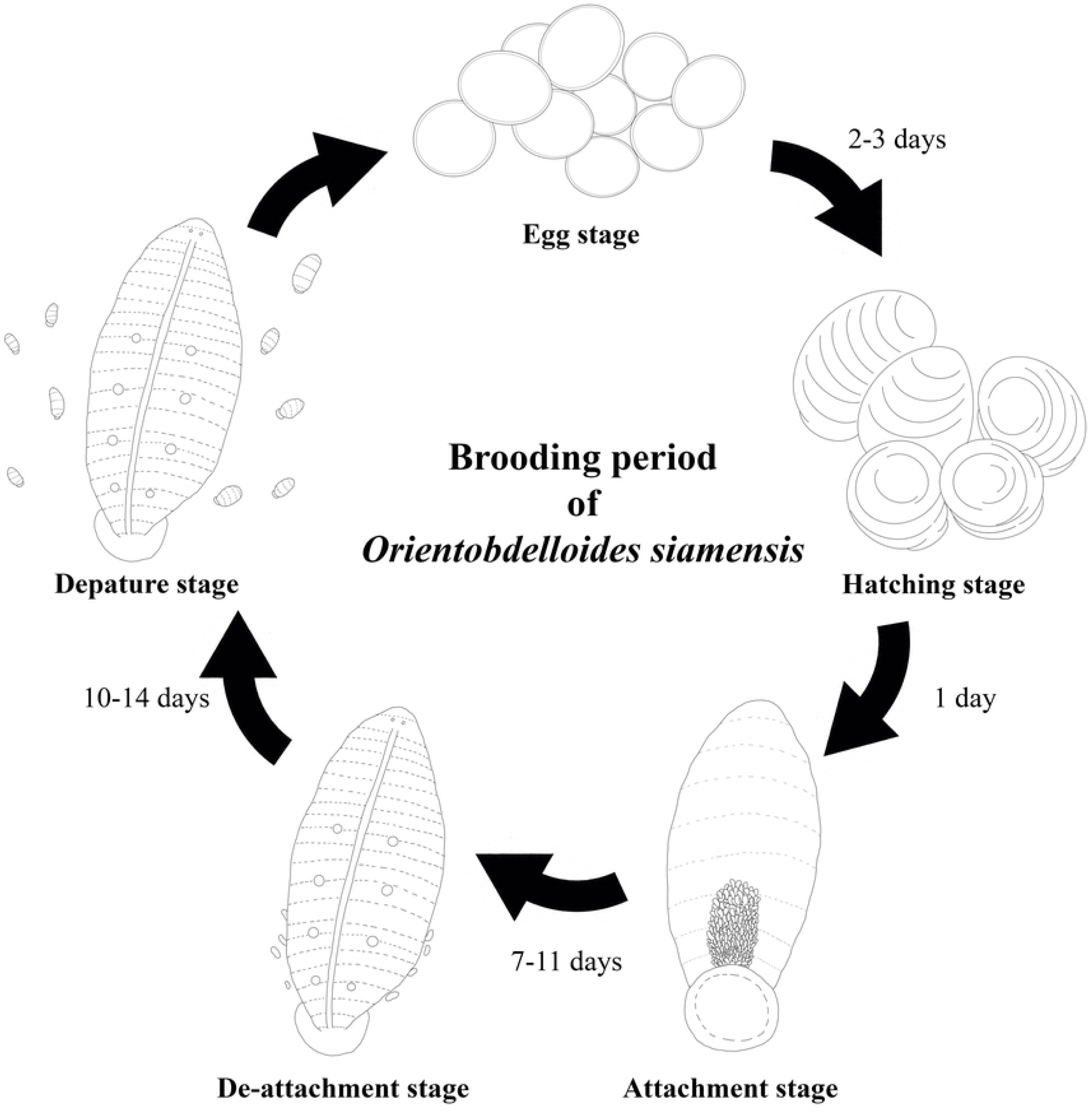
Brooding period of Orientobdelloides siamensis.

### Declaration of generative AI and AI-assisted technologies in the writing process

During the preparation of this work the authors used ChatGPT in order to Grammarly check and improve language. After using this tool/service, the authors reviewed and edited the content as needed and take full responsibility for the content of the publication.

## Acknowledgement

This research project was supported by Chulabhorn Royal Academy, Biodiversity Center from Kasetsart University, and Lee Kong Chian Natural History Museum from National University of Singapore for funding. Additionally, all tools and facilities were supported by Department of Zoology, Faculty of Science, Kasetsart University.

## Supporting information

**Supplementary Table 1.** Measurements of various characteristics in *Orientobdelloides siamensis* juveniles from hatching to parental departure (Day 1-21)

| Date | Length (L) (μm)<br>(min-max) | Width (W) (μm)<br>(min-max) | L/W | Oval area<br>(sqμm)<br>(3/4 Parent<br>area/<br>Juvenile area)<br>(individuals) | Oral sucker<br>diameter (OSD)<br>(μm)<br>(min-max) | Caudal sucker<br>diameter (CSD)<br>(μm)<br>(min-max) | CSD/<br>OSD | Yolk<br>score | Departure<br>count<br>(individual)<br>(min-max) |
| --- | --- | --- | --- | --- | --- | --- | --- | --- | --- |
| Day 0<br>(eggs) | 455.75±36.53<br>(363.28-564.97) | 409.33±39.12<br>(298.46-478.14) | 1.11 | - | - | - |  | - | - |
| Day 1<br>(hatching) | 563.30±70.76<br>(411.94-678.34) | 375.53±41.73<br>(305.87-473.61) | 1.50 | 166,205.72<br>(5,050.37) | 175.78±24.64<br>(145.54-224.47) | 246.87±23.41<br>(197.34-278.74) | 1.40 | 10.00 | 0 |
| Day 2 | 580.81±66.46<br>(483.47-821.41) | 383.13±28.69<br>(330.54-444.01) | 1.52 | 174,840.36<br>(4,800.95) | 181.15±24.69<br>(145.54-246.67) | 256.14±16.86<br>(219.54-16.86) | 1.41 | 10.00 | 0 |
| Day 3 | 614.02±66.43<br>(454.05-770.27) | 422.86±30.01<br>(348.72-471.79) | 1.45 | 204,007.65<br>(4,114.55) | 218.28±36.65<br>(148.00-294.59) | 290.55±21.76<br>(256.54-351.28) | 1.33 | 10.00 | 0 |
| Day 4 | 683.59±74.99<br>(540.54-917.14) | 459.47±74.99<br>(391.89-548.57) | 1.49 | 246,786.09<br>(3,401.33) | 217.54±35.11<br>(153.85-271.79) | 342.31±13.85<br>(305.13-371.79) | 1.57 | 9.94 | 0 |
| Day 5 | 784.81±76.39<br>(638.89-1,025.42) | 505.14±43.43<br>(443.06-586.67) | 1.55 | 311,490.91<br>(2,694.78) | 247.82±39.99<br>(175.26-309.63) | 404.01±16.34<br>(360.13-438.82) | 1.63 | 9.94 | 0 |
| Day 6 | 834.61±159.99<br>(560.00-1,273.33) | 503.76±71.64<br>(382.86-703.33) | 1.66 | 330,349.24<br>(2,540.95) | 278.68±35.56<br>(153.85-345.95) | 366.25±32.67<br>(287.18-427.03) | 1.31 | 9.74 | 0 |
| Day 7 | 1,059.60±130.22<br>(853.33-1,503.33) | 576.47±77.43<br>(463.33-806.67) | 1.84 | 479,933.21<br>(1,748.99) | 314.59±29.98<br>(232.43-362.16) | 408.86±57.27<br>(305.41-551.35) | 1.30 | 8.94 | 0 |
| Day 8 | 1,327.21±143.39<br>(1,056.00-1,804.00) | 720.62±108.96<br>(568.00-1,052.17) | 1.84 | 751,471.16<br>(1,117.01) | 409.85±52.60<br>(286.67-496.30) | 526.16±81.13<br>(376.67-722.22) | 1.28 | 8.24 | 0 |
| Day 9 | 1,672.45±330.87<br>(1,148.83-2,402.09) | 886.78±105.48<br>(673.08-1,165.87) | 1.89 | 1,165,290.58<br>(720.34) | 480.08±86.40<br>(300.75-639.10) | 613.40±118.08<br>(430.00-920.00) | 1.28 | 7.52 | 0 |
| Day 10 | 2,703.48±746.09<br>(1,860.47-5,977.44) | 1,457.04±242.73<br>(1,052.63-2,361.11) | 1.86 | 3,094,991.49<br>(271.21) | 559.69±116.33<br>(336.54-740.74) | 683.57±123.63<br>(516.83-1,045.67) | 1.22 | 6.22 | 0 |
| Day 11 | 3,480.97±564.42<br>(2,573.73-4,932.98) | 1,778.29±235.38<br>(1,293.30-2,355.66) | 1.96 | 4,863,704.19<br>(172.58) | 737.95±158.15<br>(481.93-963.86) | 936.71±162.51<br>(696.20-1,376.58) | 1.27 | 5.20 | 0 |
| Day 12 | 3,651.95±487.64<br>(2,646.10-5,016.23) | 1,454.00±209.24<br>(1,040.00-2,000.00) | 2.51 | 4,172,089.80<br>(201.19) | 872.57±181.71<br>(514.29-1,371.43) | 1,183.46±183.71<br>(808.27-1,766.92) | 1.36 | 4.40 | 0 |
| Day 13 | 4,274.81±625.46<br>(3,251.88-5,808.27) | 1,789.95±278.36<br>(1,244.02-2,392.34) | 2.39 | 6,012,057.07<br>(139.62) | 932.57±228.77<br>(571.43-1,628.57) | 1,344.84±259.88<br>(757.89-1,978.95) | 1.44 | 3.22 | 0 |
| Day 14 | 5,510.72±749.83<br>(4,164.59-8,129.68) | 2,133.33±219.91<br>(1,542.29-2,935.32) | 2.58 | 9,237,021.73<br>(90.87) | 964.41±139.52<br>(711.74-1,281.14) | 1,558.01±224.43<br>(1,103.20-2,206.41) | 1.62 | 2.08 | 0.10±0.32<br>(0-1) |
| Day 15 | 6,608.80±759.02<br>(5,190.62-7,829.91) | 2,391.01±377.23<br>(1,741.57-3,146.07) | 2.76 | 12,415,628.85<br>(67.61) | 1,008.86±153.32<br>(738.01-1,328.41) | 1,665.90±305.19<br>(1,187.74-2,375.48) | 1.65 | 1.32 | 1.20 ±1.03<br>(0-3) |
| Day 16 | 7,600.12±872.87<br>(5,969.21-9,004.40) | 2,379.77±203.26<br>(1,622.09-2,459.30) | 3.19 | 14,210,830.63<br>(59.07) | 953.72±144.94<br>(697.67-1,255.81) | 2,548.97±254.04<br>(1,603.45-2,741.38) | 2.67 | 0.60 | 2.20±0.63<br>(1-3) |
| Day 17 | 8,074.03±1,365.47<br>(5,888.11-12,086.12) | 2,623.55±520.09<br>(1,693.70-3,839.06) | 3.08 | 16,643,481.69<br>(50.43) | 956.21±270.18<br>(480.16-1,975.52) | 1,712.25±291.22<br>(1,137.83-2,675.78) | 1.79 | 0.66 | 5.00 ±0.82<br>(4-6) |
| Day 18 | 8,392.09±1,419.26<br>(6,120.06-12,562.23) | 2,721.52±539.51<br>(1,756.95-3,982.43) | 3.08 | 17,945,143.44<br>(46.78) | 1,020.50±238.34<br>(512.45-2,108.35) | 2,004.98±341.02<br>(1,332.36-3,133.24) | 1.96 | 0.42 | 5.80±1.32<br>(4-8) |
| Day 19 | 9,806.43±1,630.34<br>(6,497.46-14,517.77) | 3,004.74±660.10<br>(1,675.13-4,602.37) | 3.26 | 23,151,660.39<br>(36.26) | 1,165.82±349.57<br>(558.38-2,436.55) | 2,164.13±438.78<br>(1,404.40-3,620.98) | 1.86 | 0.14 | 13.80±0.79<br>(13-15) |
| Day 20 | 10,741.71±1,623.25<br>(6,994.54-15,628.42) | 3,281.24±676.93<br>(1,803.28-4,954.46) | 3.27 | 27,693,380.69<br>(30.31) | 1,269.58±371.50<br>(1,511.84-3,898.00) | 2,363.21±455.44<br>(1,511.84-3,898.00) | 1.86 | 0.04 | 21.30±2.79<br>(19-29) |
| Day 21 | 11,676.24±1,764.47<br>(7,603.06-16,988.09) | 3,351.13±691.35<br>(1,841.69-5,059.99) | 3.48 | 30,743,892.43<br>(30.31) | 1,269.58±371.50<br>(1,511.84-3,898.00) | 2,363.21±455.44<br>(1,511.84-3,898.00) | 1.91 | 0.00 | 135.1±35.43<br>(91-190) |
| Parent | 53,802.04±5,632.71<br>(49,232.57 -61,045.56) | 26,475.53 ± 4,609.98<br>(21,218.91- 32,175.33) | 2.03 | Full area:<br>1,119,200,760.57<br>¾ area:<br>839,400,570.42 | 9,147.65±2,715.93<br>(6,169.15-12,542.45) | 10,874.05±2,222.81<br>(8,905.47-13,418.16) | 1.19 | - | - |

**Supplementary Table 2.** Characteristics and cross-sectional area of each annuli (1-76) from *Orientobdelloides siamensis*.

| Annuli | Anterior height (μm) | Width (μm) | Posterior height (μm) | Cross-sectional area (CSA) (μm) |
| --- | --- | --- | --- | --- |
| 1 | 29.70±3.86 | 143.14±17.13 | 32.86±3.93 | 205.70±24.86 |
| 2 | 31.91±4.14 | 229.88±27.51 | 35.30±4.22 | 297.10±35.81 |
| 3 | 28.70±3.73 | 197.35±23.62 | 31.75±3.80 | 257.80±31.08 |
| 4 | 44.96±5.84 | 223.38±26.74 | 49.73±5.95 | 318.06±38.43 |
| 5 | 40.34±5.24 | 130.12±15.57 | 44.62±5.34 | 215.09±26.07 |
| 6 | 46.76±6.07 | 130.12±15.57 | 51.73±6.19 | 228.61±27.75 |
| 7 | 51.98±6.75 | 114.94±13.76 | 57.50±6.88 | 224.42±27.29 |
| 8 | 40.14±5.21 | 117.11±14.02 | 44.40±5.31 | 201.65±24.46 |
| 9 | 42.35±5.50 | 91.09±10.90 | 46.84±5.61 | 180.28±21.93 |
| 10 | 48.77±6.33 | 88.92±10.64 | 53.95±6.46 | 191.64±23.34 |
| 11 | 50.38±6.54 | 114.94±13.76 | 55.72±6.67 | 221.04±26.87 |
| 12 | 64.63±8.39 | 88.92±10.64 | 71.48±8.56 | 225.03±27.48 |
| 13 | 51.78±6.72 | 132.29±15.83 | 57.28±6.86 | 241.35±29.31 |
| 14 | 69.24±8.99 | 140.97±16.87 | 76.59±9.17 | 286.80±34.90 |
| 15 | 70.45±9.15 | 117.11±14.02 | 77.92±9.33 | 265.48±32.36 |
| 16 | 76.67±9.95 | 130.12±15.57 | 84.80±10.15 | 291.60±35.54 |
| 17 | 74.46±9.67 | 88.92±10.64 | 82.36±9.86 | 245.74±30.04 |
| 18 | 70.85±9.20 | 104.10±12.46 | 78.37±9.38 | 253.31±30.91 |
| 19 | 79.28±10.29 | 104.10±12.46 | 87.69±10.50 | 271.07±33.11 |
| 20 | 129.46±16.81 | 127.95±15.31 | 143.19±17.14 | 400.60±49.05 |
| 21 | 126.85±16.47 | 140.97±16.87 | 140.30±16.79 | 408.12±49.92 |
| 22 | 130.46±16.94 | 151.81±18.17 | 144.30±17.27 | 426.57±52.16 |
| 23 | 99.35±12.90 | 140.97±16.87 | 109.89±13.15 | 350.21±42.75 |
| 24 | 71.25±9.25 | 114.94±13.76 | 78.81±9.43 | 265.00±32.31 |
| 25 | 84.30±10.94 | 91.09±10.90 | 93.24±11.16 | 268.62±32.87 |
| 26 | 85.50±11.10 | 153.98±18.43 | 94.57±11.32 | 334.05±40.69 |
| 27 | 86.10±11.18 | 177.84±21.28 | 95.24±11.40 | 359.18±43.70 |
| 28 | 84.70±11.00 | 203.86±24.40 | 93.68±11.21 | 382.24±46.45 |
| 29 | 76.67±9.95 | 206.03±24.66 | 84.80±10.15 | 367.50±44.61 |
| 30 | 114.80±14.91 | 190.85±22.84 | 126.98±15.20 | 432.64±52.75 |
| 31 | 110.19±14.31 | 153.98±18.43 | 121.88±14.59 | 386.04±41.13 |
| 32 | 89.51±11.62 | 140.97±16.87 | 99.01±11.85 | 329.49±40.19 |
| 33 | 96.34±12.51 | 164.82±19.73 | 106.56±12.75 | 367.72±44.82 |
| 34 | 96.74±12.56 | 242.90±29.07 | 107.00±12.81 | 446.64±54.25 |
| 35 | 104.37±13.55 | 268.92±32.19 | 115.44±13.82 | 488.73±59.35 |
| 36 | 103.97±13.50 | 292.78±35.04 | 115.00±13.76 | 511.74±62.10 |
| 37 | 83.90±10.89 | 307.96±36.86 | 92.80±11.11 | 484.65±58.69 |
| 38 | 70.65±9.17 | 318.80±38.16 | 78.14±9.35 | 467.59±56.53 |
| 39 | 94.73±12.30 | 333.98±39.97 | 104.78±12.54 | 533.50±64.62 |
| 40 | 105.97±13.76 | 383.86±45.94 | 117.22±14.03 | 607.05±73.52 |
| 41 | 102.36±13.29 | 268.92±32.19 | 113.22±13.55 | 484.50±58.83 |
| 42 | 95.13±12.35 | 206.03±24.66 | 105.23±12.59 | 406.39±49.43 |
| 43 | 128.45±16.68 | 153.98±18.43 | 142.08±17.01 | 424.51±51.89 |
| 44 | 84.10±10.92 | 127.95±15.31 | 93.02±11.13 | 305.07±37.22 |
| 45 | 85.50±11.10 | 229.88±27.51 | 94.57±11.32 | 409.96±49.77 |
| 46 | 73.06±9.49 | 357.84±42.83 | 80.81±9.67 | 511.70±61.83 |
| 47 | 87.91±11.41 | 242.90±29.07 | 97.24±11.64 | 428.04±51.95 |
| 48 | 106.78±13.86 | 190.85±22.84 | 118.10±14.14 | 415.73±50.65 |

| **Annuli** | **Anterior height**<br>**($\mu\text{m}$)** | **Width**<br>**($\mu\text{m}$)** | **Posterior height**<br>**($\mu\text{m}$)** | **Cross-sectional**<br>**area (CSA) ($\mu\text{m}$)** |
| --- | --- | --- | --- | --- |
| 49 | 89.72±11.65 | 318.80±38.16 | 99.23±11.88 | 507.75±61.50 |
| 50 | 130.46±16.94 | 294.95±35.30 | 144.30±17.27 | 569.71±69.26 |
| 51 | 126.85±16.47 | 344.83±41.27 | 140.30±16.79 | 611.98±74.29 |
| 52 | 93.93±12.20 | 279.77±33.48 | 103.90±12.43 | 477.59±57.93 |
| 53 | 51.38±6.67 | 273.26±32.71 | 56.83±6.80 | 381.47±46.07 |
| 54 | 33.64±4.37 | 362.18±43.35 | 37.21±4.45 | 433.02±52.09 |
| 55 | 34.20±4.44 | 347.00±41.53 | 37.83±4.53 | 419.03±50.42 |
| 56 | 29.22±3.79 | 318.80±38.16 | 32.32±3.87 | 380.35±45.75 |
| 57 | 35.16±4.57 | 320.97±38.42 | 38.89±4.66 | 395.03±47.56 |
| 58 | 42.71±5.55 | 357.84±42.83 | 47.24±5.65 | 447.79±53.93 |
| 59 | 35.89±4.66 | 271.09±32.45 | 39.69±4.75 | 346.67±41.78 |
| 60 | 52.18±6.78 | 320.97±38.42 | 57.72±6.91 | 430.87±51.99 |
| 61 | 50.74±6.59 | 323.14±38.68 | 56.12±6.72 | 430.00±51.87 |
| 62 | 93.93±12.20 | 258.08±30.89 | 103.90±12.43 | 455.90±55.34 |
| 63 | 99.00±12.85 | 290.61±34.78 | 109.51±13.11 | 499.12±60.55 |
| 64 | 88.53±11.49 | 234.22±28.03 | 97.92±11.72 | 420.67±51.08 |
| 65 | 94.07±12.21 | 210.37±25.18 | 104.05±12.45 | 408.49±49.67 |
| 66 | 88.12±11.44 | 229.88±27.51 | 97.46±11.67 | 415.46±50.45 |
| 67 | 88.73±11.52 | 173.50±20.77 | 98.15±11.75 | 360.38±43.87 |
| 68 | 103.73±13.47 | 208.20±24.92 | 114.73±13.73 | 426.65±51.93 |
| 69 | 98.59±12.80 | 199.52±23.88 | 109.05±13.05 | 407.16±49.55 |
| 70 | 95.30±12.37 | 177.84±21.28 | 105.42±12.62 | 378.56±46.10 |
| 71 | 71.07±9.23 | 273.26±32.71 | 78.61±9.41 | 422.93±51.20 |
| 72 | 66.14±8.59 | 251.57±30.11 | 73.15±8.76 | 390.87±47.32 |
| 73 | 54.43±7.07 | 290.61±34.78 | 60.21±7.21 | 405.24±48.94 |
| 74 | 50.94±6.61 | 314.47±37.64 | 56.34±6.74 | 421.75±50.88 |
| 75 | 38.20±4.96 | 299.28±35.82 | 42.26±5.06 | 379.75±45.75 |
| 76 | 47.65±6.19 | 180.00±21.54 | 52.71±6.31 | 280.36±33.94 |
| Average of anterior<br>part (annuli 1-29) | 68.55±28.25<br>(28.70-130.46) | 137.83±40.98<br>(88.92-229.88) | 75.82±31.25<br>(31.75-144.30) | 282.20±69.90<br>(180.28-426.57) |
| Average of<br>posterior part<br>(annuli 30-76) | 81.25±27.63<br>(29.22-130.46) | 261.77±67.08<br>(127.95-383.86) | 89.87±30.56<br>(32.32-144.30) | 432.90±68.31<br>(280.36-611.98) |
| Total average<br>(annuli 1-76) | 84.51±31.38<br>(31.75-144.30) | 241.48±84.03<br>(88.92-383.86) | 76.41±28.37<br>(28.70-130.46) | 375.40±100.28<br>(180.28-611.98) |

